# NuclePhaser: a YOLO-based framework for cell nuclei detection and counting in phase contrast images of arbitrary size with support of fast calibration and testing on specific use cases

**DOI:** 10.1101/2025.05.13.653705

**Authors:** Nikita Voloshin, Egor Putlyaev, Elizaveta Chechekhina, Vladimir Usachev, Maxim Karagyaur, Kirill Bozov, Olga Grigorieva, Pyotr Tyurin-Kuzmin, Konstantin Kulebyakin

**Affiliations:** Department of Biochemistry and Regenerative Biomedicine, Faculty of Medicine, Medical Research and Educational Institute, Moscow State University, 119192, Moscow, Russia; Laboratory of Physical Сhemistry of Analytical Processes, N. M. Emanuel Institute of Biochemical Physics, Russian Academy of Sciences, 119334, Moscow, Russia; Center for Regenerative Medicine, Medical Research and Educational Institute, Moscow State University, 119192, Moscow, Russia

## Abstract

Microscopy is an essential method in modern biology, and brightfield microscopy methods (phase contrast, differential interference contrast, etc.) are being widely used and actively developed since they don’t require sample fixation and staining. But they produce low contrast images, where cells have similar intensity to the background. In this work we developed and tested a set of deep learning object detection YOLO models that detect cell nuclei in phase contrast images, which allows cell count and tracking without staining. We created a large dataset consisting of more than 100,000 640×640 pixels images with more than 3 million nuclei of 4 different cell cultures (CHO, HEK293, iPSCs, and MSCs). Using images from various microscopes and cameras, as well as full-scale augmentations, we developed a set of highly generalized models that can detect nuclei in images across different cell types and imaging conditions, including different microscopes and contrast methods. Combined with sliced inference methods, these algorithms can be applied to images of any size, allowing studies of large quantities of cells. Moreover, we developed a training-free calibration and testing algorithm based on confidence threshold optimization. It allows for fine-tuning of models for specific cell types and/or imaging options and evaluating the accuracy of the calibrated model. This provides a highly controllable and reliable method for studying cell proliferation rate, single cell tracking and other scenarios. Additionally, we developed a NuclePhaser plugin for Napari (https://github.com/nikvo1/napari-nuclephaser), which allows users to calibrate, test and apply our models in code-free manner. Given that the YOLO models are fast and can run at sufficient speeds even on CPUs, this makes our work highly accessible to a wide range of researchers.

## Introduction

Microscopy is a fundamental method in medical and biological sciences. There are many different microscopic techniques, and each of them has a unique set of advantages and drawbacks [1]. However, the majority of microscopy methods prevent studying dynamics of cell cultures: they require fixation, which stops all cellular processes, and/or staining, which can also interfere with cell physiology [2].

Brightfield microscopy methods, on the other hand, don’t require fixation or staining, and are widely used in studying cell processes in dynamics. Despite this, they pose challenges regarding automatic image processing due to the similar intensity of cell pixels and the background, even with use of contrast enhancement methods such as phase contrast or differential interference contrast (DIC). There are several studies in which traditional computer vision techniques were applied to cell detection and counting in brightfield images [3–5], but creation of algorithms generalized for different complex cell shapes and imaging conditions remains a challenge.

Several deep learning methods address this challenge, including various image-to-image translation [6,7], whole-cell segmentation [8,9], and nuclei segmentation [10,11].

The alternative approach we propose in this work is a fast and generalized to various cell types and imaging conditions nuclei detection model based on YOLO algorithms. Key features of our approach are an ability to detect nuclei on large whole-slide images of arbitrary sizes with Slicing Aided Hyper Inference [12]; an algorithm to calibrate confidence threshold for exact images and test calibrated model accuracy at counting nuclei; and an open-source NuclePhaser plugin for Napari, that allows users to perform inference, count nuclei, convert detections to labels suitable for tracking, and calibrate and test models against various ground truths without coding.

There are a lot of scenarios where nuclei count or localization on a brightfield image could be beneficial. But here we highlight two areas where our system offers advantages over existing methods: assessing cell population growth rate and single-cell tracking. Studying cell growth rate is widely used in cancer studies, drug development and other areas [13]. Various methods of cell population growth assessment exist, but some of them don’t directly measure the number of cells, but rather derive it from cells metabolic activity (for example, MTT assay [13]). Others estimate the area of substrate occupied by the population [14], which means that spreading or contraction of individual cells can imitate growth rate changes. Current methods that explicitly count cells on time-lapse microscopic images either require fluorescent dyes, or rely on the basis of the instance segmentation approach [15], which has certain limitations regarding inference on large whole-slide images and computational demands. Here we propose a highly generalized and fast alternative for assessing cell growth by counting the number of nuclei on time-lapse brightfield images. We also provide simple pipelines for calibrating and testing models for specific sets of images, so users can be confident knowing the accuracy of the method on their specific use cases.

Single cell tracking on brightfield images is another problem where our nuclei detection models can be useful. It’s a challenging task as it requires consistent generation of labels in a set of consequent frames with single error spoiling the whole track. We tested our models on publicly available Cell Tracking Challenge datasets [16] as generators of labels for single cell tracking. Our approach offers high generalization, fast inference, ability to work on arbitrary sizes of images, and convenience to use with tracking algorithms distributed as Napari plugins (in particular, btrack [17]), since inference on a timelapse array of images is included as an option in our NuclePhaser plugin. Combined with Napari’s set of tools for manual correction of detections, it can also be used as a fast and convenient semiautomated alternative for this task.

We used Ultralytics YOLOv5 and YOLOv11 families of object detection models. To achieve high generalization of models, we used a combination of approaches on different levels of model creation. First of all, we used different cell lines, microscopes, magnifications and cameras for dataset creation. Secondly, we used an auxiliary object detection model that detects nuclei on DAPI images for automatic labeling of images, which allowed us to create a large dataset with over 100,000 images and more than 3 million nuclei. Third, we used a set of full-scale augmentations [18] during model training, artificially increasing dataset size and imitating different microscopy settings and artifacts (illumination, magnification, defocus, etc.). Our models demonstrated high generalization, demonstrating the ability to count nuclei with high accuracy on the LIVE-Cell dataset [15], which consists of images of different cell types taken with a different microscope system.

## Results

### Auxiliary fluorescent nuclei detector

For automatic labeling of nuclei in phase contrast images, a series of fluorescent nuclei detection models was trained. We collected a small dataset consisting of 191 manually annotated DAPI images, which were divided into 156 training images and 35 validation images, each with a resolution of 640 x 640 pixels. Using this dataset, we trained YOLOv5 and YOLOv11 models. The results are presented in fig. 1. mAP 0.5 of these models surpasses 0.95 even for smallest ones, which is expected, since detecting bright circles on black background is a relatively simple computer vision task.

**Figure 1.**
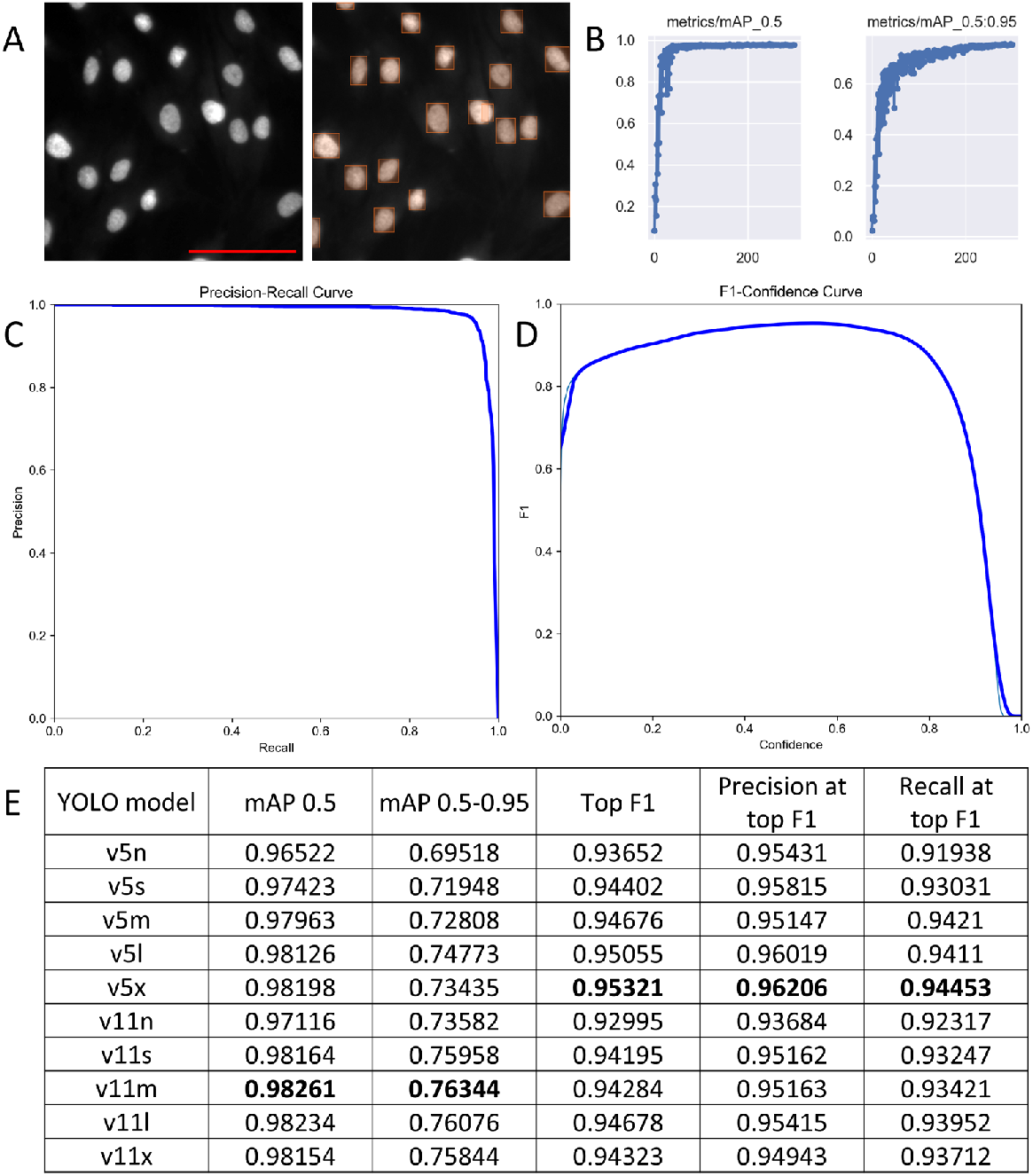
Training auxiliary fluorescent nuclei detection models. **(A)** An example image from the training dataset (left) and its manually annotated version (right), screenshot from makesense.ai, scale bar (red): 100 μm. **(B)** Changes of validation mAP 0.5 and mAP 0.5-0.95 during training of the YOLOv5x model for 250 epochs. **(C)** Precision-recall curve of YOLOv5x model on validation dataset. **(D)** F1-confidence threshold curve of YOLOv5x model for validation dataset. **(E)** Table of validation results for YOLOv5 and YOLOv11 models on the fluorescent nuclei dataset, best results are highlighted in bold.

### Phase contrast images dataset creation

We used fluorescent nuclei detection models for automatic labeling of a large dataset of phase contrast images (see fig. 2). The YOLOv5x fluorescent nuclei detection model was used with a confidence threshold of 0.5. The resulting dataset contains a total of 112,032 images (divided into 90987 train and 21045 validation) and 3,997,168 nuclei, distribution of which among different cell types is presented in fig. 2. To find out the automatic labeling error, we randomly sampled 100 images from the dataset and manually checked for false negative and false positive labels. Among 6054 nuclei on 100 images there were 5 (0.083%) false positives and 110 (1.817%) false negatives (majority is in the form of two closely located nuclei detected as one merged object, see fig. 2). All nuclei were labeled with the same class “Nucleus”, i.e. the result dataset is single-class, since there is no task of classifying the type of cell on an image by detected nuclei.

**Figure 2.**
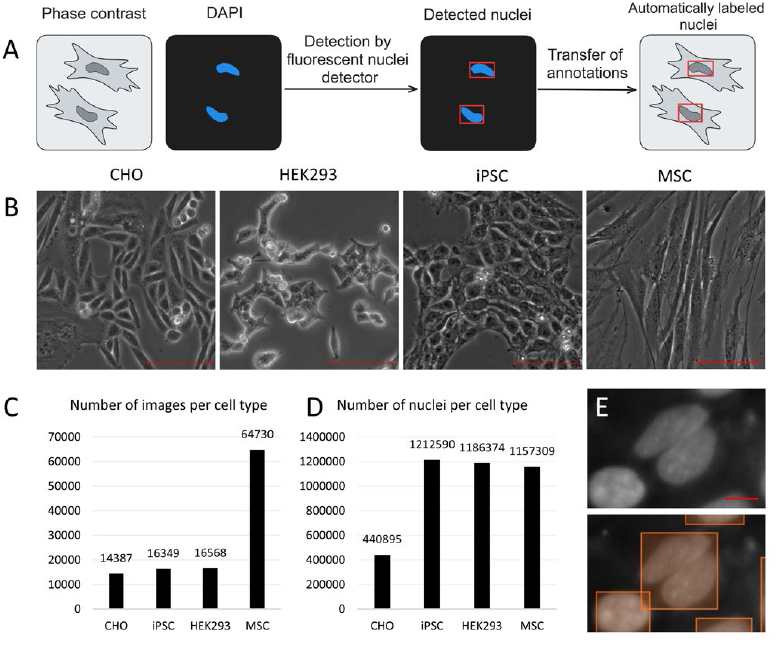
Creating large phase contrast images dataset with automatic labeling by fluorescent nuclei detection model. 4**(A)** Workflow diagram of automatic labelling. **(B)** Examples of phase contrast images of each cell type, scale bars: 100 μm. **(C)** Distribution of number of images among four cell types. **(D)** Distribution of number of nuclei on all images of four cell types. **(E)** Example of automatic labeling error, false negative detection in the form of two closely located nuclei merged into one object, scale bar: 10 μm.

### Models training

Using this large dataset, we trained a set of YOLOv5 and YOLOv11 phase contrast nuclei detection models; validation results are presented in fig. 3. We used a set of augmentations from Albumentations library [18] to emulate different microscopy settings and artifacts. Models were trained with different schedules, but all until peak or plato on validation mAP 0.5. Largest mAP 0.5 of 0.878, as well as top F1, precision at top F1 and recall at top F1, were reached by the YOLOv5l model (see fig. 3). To evaluate models performance on four cell types in the validation dataset, we conducted a set of validations on subsets with separate cell types, with results detailed in Supplementary table 1.

**Figure 3.**
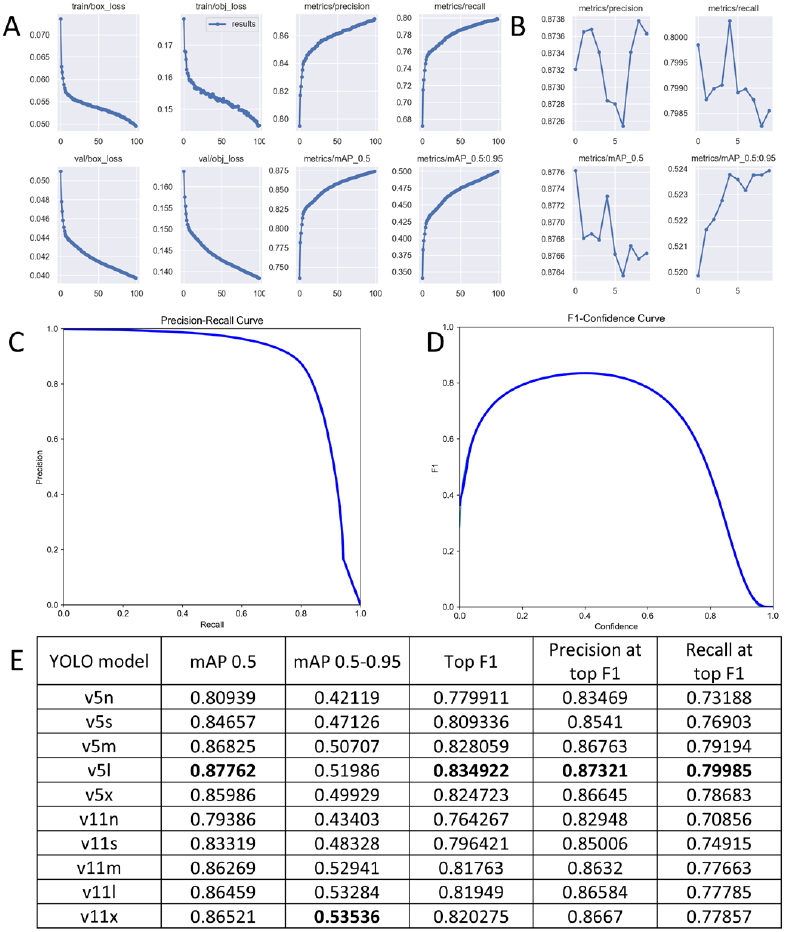
Training phase contrast nuclei detection YOLO models. **(A)** Graphs representing training progress of the YOLOv5l model for 100 epochs during the first stage. **(B)** Graphs representing training progress of the same YOLOv5l model for 10 epochs during the second stage with lower learning rate. **(C)** Precision-recall curve for YOLOv5l model on validation dataset. **(D)** F1-confidence threshold curve for YOLOv5l model on validation dataset. **(E)** Table of validation results of YOLOv5 and YOLOv11 models on phase contrast images with automatically annotated nuclei, with the best results highlighted in bold.

### Test of counting on LIVECell

Our first biologically oriented test assessed models’ performance in the cell counting. We used the LIVECell dataset – a collection of phase contrast images of 8 cell types, none of which was in the training or validation datasets for our models. In addition, these images were obtained with a different microscope system. Since the number of objects detected by the model is highly dependent on the model’s confidence threshold hyperparameter, we used a part of the dataset to calibrate it, iteratively selecting the confidence threshold returning lowest counting error (see fig. 4). For each of the 8 cell types we combined train, validation and test parts into one dataset, 10% of which was used for calibration and 90% for test; results for best performing models are presented in fig. 4, results for all models can be found in supplementary table 2.

**Figure 4.**
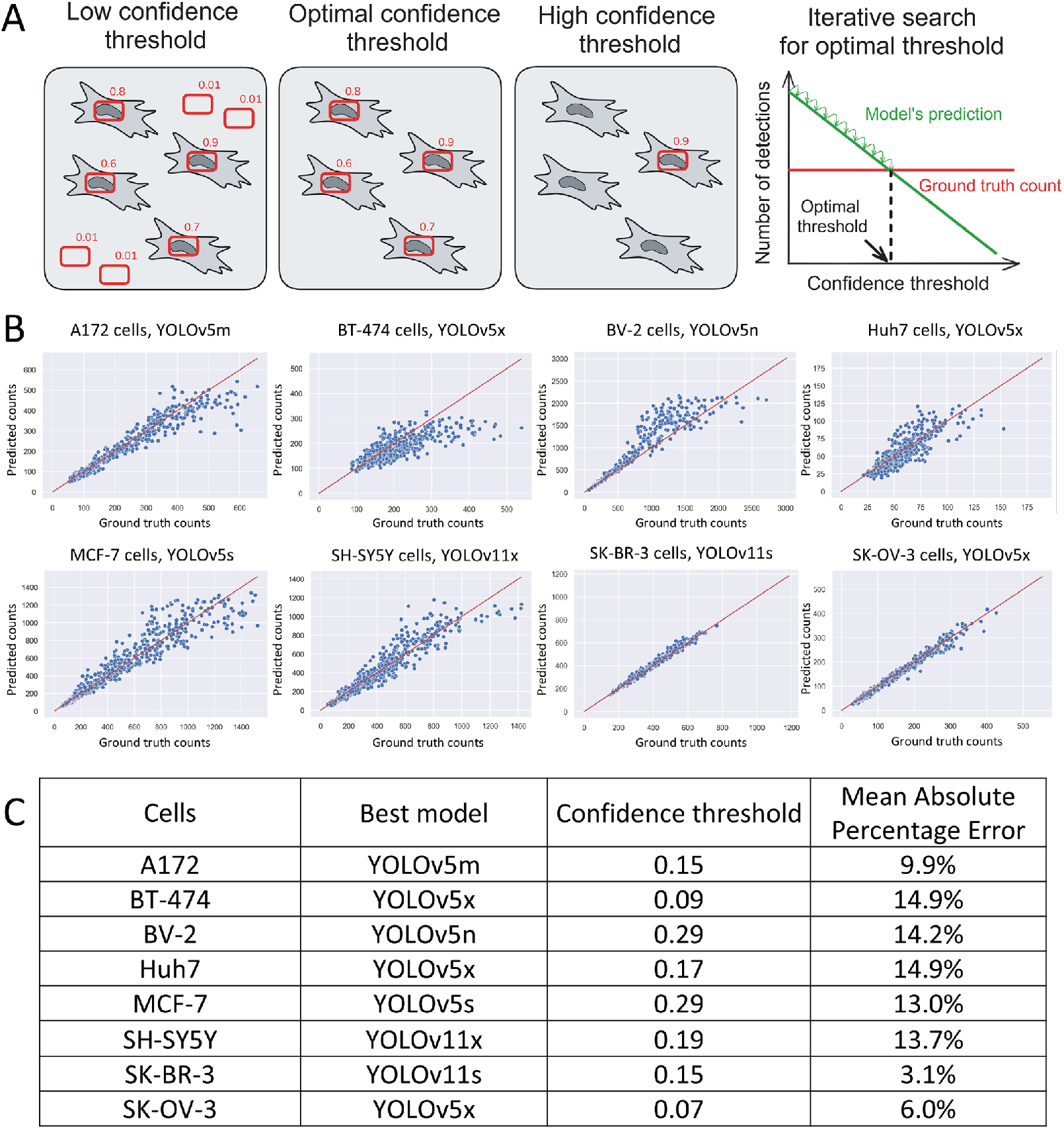
Results of counting test on LIVECell dataset. **(A)** Graphic explanation of calibration principle. **(B)** Predicted vs. Ground truth counts scatterplots for each of 8 cell types (best YOLO models are shown). Every point represents an image. Red line is the imaginary line of perfect predictions. **(C)** Table of best-performing models and their corresponding results for each cell type.

Despite the fact that models didn’t train on any of the 8 cell types of LIVECell, they perform well in counting these cells, especially in images of SK-BR-3 and SK-OV-3 cells with MAPE of 3.1% and 6.0% respectively. Unexpectedly, for some cells the best performing models are small ones, for example, the YOLOv5n model performs best on BV-2 cells and YOLOv5s – on MCF-7.

### Test of tracking on Cell Tracking Challenge datasets

The second biologically oriented test we performed was assessing whether our models predictions are suitable for single cell tracking in consecutive frames. We used Cell Tracking Challenge datasets for that: U373 dataset to test how models perform on phase contrast images, and HeLa dataset to test how models will perform on differential interference contrast, a different imaging modality. Btrack [17] plugin was used as a tracker, and tracks derived from manual annotations were used as ground truth. Two biology-motivated metrics were used: Complete Tracks (CT), which evaluates how well the tracking system suits the task of end-to-end tracking; and Branching Correctness (BC), which evaluated how accurate the system is on prediction cell divisions. Results are presented in fig. 5.

**Figure 5.**
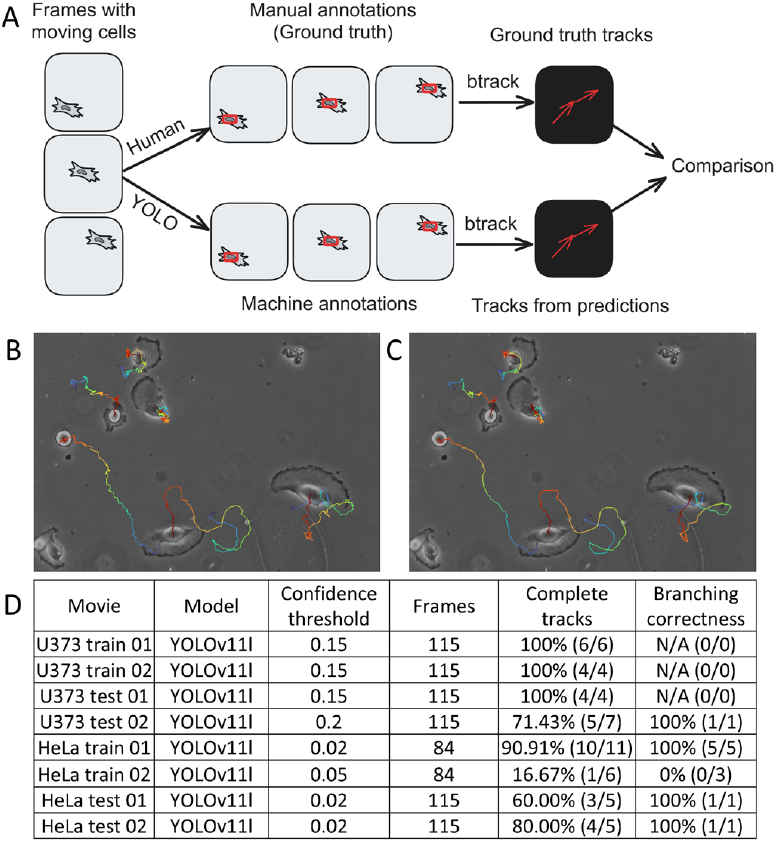
Test of model’s predictions fitness as tracking markers generator. **(A)** Workflow diagram of the test pipeline. **(B)** Final frame of U373 train 01 movie with end-to-end tracks reconstructed from manual annotations, screenshot from Napari, different colors on tracks represent different time points. **(C)** Same frame with end-to-end tracks reconstructed from annotations made by YOLOv11l model, screenshot from Napari. **(D)** Table of tracking test results on 8 movies from Cell Tracking Challenge, Complete tracks and Branching correctness metrics are derived from comparison of machine annotations-based tracks to manual annotations-based ones.

### Napari plugin

To increase availability of our work, we developed the NuclePhaser plugin to Napari (see fig. 6). NuclePhaser is available as an open-source repository on GitHub (https://github.com/nikvo1/napari-nuclephaser). The functionality of NuclePhaser plugin includes:

**Figure 6.**
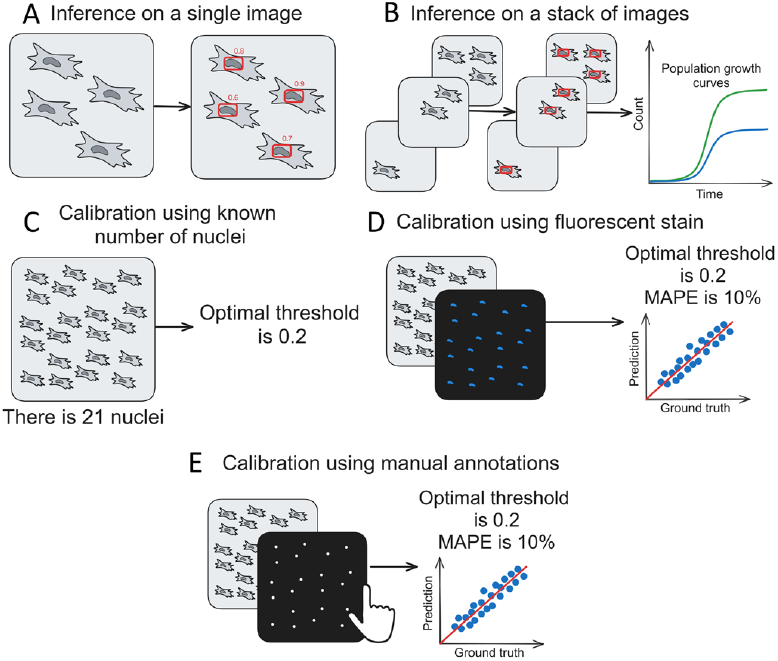
Graphic representations of NuclePhaser Napari plugin functionality. **(A)** Plugin allows inference with YOLO and SAHI on single images. **(B)** Plugin allows inference on stacks of images, automatically producing a table in .csv or .xlsx format with counting results for each frame, which can be used to reconstruct population growth curves (graph drawing is not included in plugin functionality). **(C)** Plugin allows optimization of a given model’s confidence threshold with known number of objects on a calibration image; this method of calibration doesn’t produce accuracy metrics. **(D)** Plugin allows calibration of a given model’s confidence threshold with a brightfield image – fluorescent stain image pair; method returns optimal threshold and accuracy metrics – MAPE and scatterplot. **(E)** Same calibration method as in D, but, instead of fluorescent image, a manual detections of nuclei is used as ground truth.

- Inference with YOLO models using SAHI sliced inference on single images or stacks of images of arbitrary sizes, including large whole-slides ones.
- Calibrating model’s confidence threshold using different ground truths:
  ○ General number of nuclei on an image without providing their localization. It’s the simplest calibration method, but it doesn’t provide accuracy metrics.
  ○ Image with fluorescent stain of nuclei. This method uses fluorescent nuclei detectors as ground truth generators. Large images can be divided into smaller slices for processing. A subset of these slices is used for calibration, while the remaining slices are reserved for testing. Plugin generates MAPE metric and a Predicted Count – Ground Truth Count scatterplot.
  ○ Manual detection of nuclei. This method follows the same principle as fluorescent stain calibration but uses manually annotated nuclei (Napari Points layer) as ground truth.
- A widget for converting detections from Points layer to Labels layer to be processed further by tracking plugins (btrack plugin, in particular, demands annotations as Labels layer)

## Discussion

One of the main obstacles of processing brightfield microscopy images is low contrast, which makes brightfield images difficult to automatically analyze. Here we present a deep learning object detection-based approach to detect nuclei on phase contrast (and, as we show, different brightfield microscopy methods as well) images with main applications for cell count and single cell tracking.

Major challenge for such a deep learning system is generalization, since it should work on a wide array of cell types, microscopes, magnifications, cameras and imaging conditions. We approached that problem on different levels. Firstly, we created a large dataset with more than 100,000 images of four cell types taken with three different cameras and with different magnifications ranging from 40x to 300x. Secondly, during training of the models we used a set of fullscale augmentations imitating various imaging conditions: contrast, focus, camera artifacts, etc. This resulted in a high level of generalization of models, which, as we show on LIVECell and Cell Tracking Challenge datasets, are capable of detecting nuclei of different cell types, taken with different microscope systems and even in images of different brightfield modality.

We tested our models in two biologically oriented scenarios. First is counting the number of nuclei on an image, which can be useful in various tasks, but it can be especially beneficial for assessing cell population growth rate by consequent images. We showed that on LIVECell dataset, consisting of images of 8 cell types that weren’t in the training dataset and taken with different microscope system, our models are capable of counting number of nuclei with MAPE ranging from 14.9% for BT-474 and Huh7 cells up to 3.1% for SK-BR-3. Combined with a calibration algorithm that produces accuracy metrics for specific images and cells, our models can be a valuable and reliable addition to a range of methods for assessing cell population growth.

Second biologically oriented scenario we tested is single cell tracking. We showed that on the Cell Tracking Challenge dataset of U373 cells, a cell line that also wasn’t in the training dataset, our models can produce stable tracking markers that, combined with btrack plugin [17], solve automatic end-to-end tracking of individual cells. We also showed that our models can produce stable tracking markers on DIC images of Hela, which are not only images of different cells taken with a different microscope system, but using a different imaging technique; although performance becomes less stable with shift to DIC.

Important feature of various computer vision methods availability is open-source publication in code-free manner [22], which expands the range of possible users. We developed a NuclePhaser plugin to Napari, functionality of which includes inference on single images or stacks of images, which makes it convenient for both cell population growth assays and single cell tracking studies. The important feature we incorporated is ability to calibrate models, i.e. adjust model’s confidence threshold for specific cell types and images, enabling rapid tuning (seconds to minutes) without modifying model weights with deep learning algorithms that demand GPU. The crucial option of calibration widgets is returning accuracy metrics – MAPE and Prediction vs. Ground Truth scatterplot. This allows fast adjusting to specific use cases along with confidence in the model’s performance, which is substantial for scientific methods.

## Conclusion

We trained a set of YOLO models to detect cell nuclei on phase contrast images. Result models are highly generalized to various cell types and microscopy systems due to the large dataset and set of full-scale augmentations. Our models demonstrated suitability for counting the number of cells on the LIVECell dataset and for producing stable tracking markers on Cell Tracking Challenge datasets. We incorporated our developments in the form of open-source NuclePhaser plugin for Napari, which makes it available for a wide range of users. The result algorithm is quickly tunable without engagement of GPU-dependent deep learning algorithms and highly reliable due to the function of calculating accuracy metrics for each specific use case.

## Code availability

NuclePhaser plugin for Napari with trained models’ weights can be found at https://github.com/nikvo1/napari-nuclephaser.

## Acknowledgements

The work was supported by the Non-commercial Foundation for Support of Science and Education “INTELLECT”.

This paper was typeset with the bioRxiv word template by @Chrelli: www.github.com/chrelli/bioRxiv-word-template

## Materials and Methods

### Cell cultures

We used four cell cultures: mesenchymal stem cells (MSCs), CHO, HEK293 and induced pluripotent stem cells (iPSCs). MSCs are represented by different groups: native primary MSCs, primary MSCs differentiated in adipogenic direction, hTERT-immortalized ASC52Telo (ATCC^®^ SCRC-4,000^TM^); these cells were obtained and cultured as previously described [19]. CHO,HEK293 and skin fibroblast-derived iPSC cells were obtained from the biobank of the Institute for Regenerative Medicine, Lomonosov MSU (https://human.depo.msu.ru). CHO, HEK293 and iPSC cells were seeded in a 6-well culture plate in a gradient of densities, starting from monolayer and decreasing density 2-fold in each consequent well. MSCs images were collected in various scenarios, dishes and densities.

### Fixation, staining and microscopy

For better preservation of cell morphology, we used a 0.2% glutaraldehyde-based fixation solution, protocol described in [20]. After fixation, cells were washed 3 times with PBS solution and stained with DAPI for 10 minutes followed by PBS washing 3-4 times.

Two microscope systems were used for acquiring images, Nikon Eclipse Ti2 with two cameras: Kinetix (Teledyne Photometrics) and Ri2 (Nikon); and Nikon Eclipse Ti with DS-Q1Mc camera. For 40x magnification Nikon Plan Fluor 4x/0.13 objective was used, for 100x – Nikon Plan Fluor 10x/0.30, for 200x – Nikon S Plan Fluor LWD 20x/0.70. 60x, 150x and 300x magnifications were achieved by using the same objectives and turning on an additional 1.5x magnification knob (physical additional lens) on both microscopes. Phase contrast and DAPI images were acquired sequentially with automatic image registration afterwards using the ND Acquisition lambda option, and the Large image option was used to acquire large fields. We captured large fields of view, resulting in images from 15000 x 15000 pixels up to 30000 x 30000 pixels.

For preprocessing, Auto Contrast function from NIS Elements GA3 module was used for both Phase and DAPI images; all images were transformed into a uniform 8-bit RGB format.

### Datasets creation

To create a dataset of annotated fluorescent nuclei, we cropped random 640×640 pixel pictures from DAPI images of 4 cell lines described above. We used makesense.ai web-based tool for manual annotations of nuclei. Phase contrast images dataset was created automatically by applying a custom slicing function to large images to get 640 x 640 pixels images and annotating them by fluorescent nuclei detection models. Datasets were split into two parts – 80% train subset and 20% validation subset. To test how models perform in the real-world scenarios, we used open-source datasets – LIVECell and Cell Tracking Challenge.

### Models train

We used Ultralytics YOLOv5 and YOLOv11 models collections for both auxiliary fluorescent nuclei detector and main phase contrast nuclei detector. To train our models, we used default train scripts provided by the developer (train.py in case of YOLOv5 and model.train() function in case of YOLOv11). NVIDIA Quadro RTX 4000 graphics processor was used for training the models. All models were trained with default training settings, with maximal batch size allowed by 8 GB of VRAM. SGD was used as an optimizer, initial learning rate was 0.01. Models were trained for different amounts of epochs and with different learning rate schedules (due to resources economy), but all were trained until peak or plato on validation mAP 0.5 metric. Plots were generated automatically by the training scripts. Albumentations library [18] augmentations were used by explicitly changing utils/augmentations.py file in case of YOLOv5 and data/augment.py in case of YOLOv11.

Augmentations used include:

- RandomRotate90(p = 0.75)
- Defocus(radius = (3, 16), p = 0.2)
- Blur(blur_limit = 11, p = 0.2)
- RandomBrightnessContrast(brightness_limit = 0.4, contrast_limit = 0.4, p = 0.2)
- GaussNoise(var_limit = (10.0, 75.0), mean = 0, per_channel = False, p = 0.2)
- PixelDropout(dropout_prob = 0.05, per_channel = False, drop_value = 0, p = 0.2)
- ImageCompression(quality_lower = 75, p = 0.2)
- RandomFog(p = 0.2)

Default accuracy metrics provided by the Ultralytics were used to validate models. Metrics used are mAP 0.5, mAP 0.5-0.95, top F1 score, precision at top F1 and recall at top F1.

- Precision is defined as the proportion of true positives among all positive predictions.
- Recall is defined as the proportion of true positives among all actual positives.
- F1-score is defined as (2 * Precision * Recall)/(Precision + Recall). Top F1 score is calculated by iterating over all confidence thresholds and searching for one returning maximal F1.
- mAP 0.5, mean Average Precision at Intersection over Union (IoU) threshold 0.5, is defined as averaged precision across all confidence thresholds across all classes (our case is single-class) with fixed IoU of 0.5.
- mAP 0.5-0.95, mean Average Precision at IoU thresholds from 0.5 to 0.95, is defined the same as mAP 0.5, but instead of fixed IoU, all mAPs across IoU thresholds from 0.5 to 0.95 with increment of 0.05 are averaged.

### Inference and tests

We used obss/sahi library [12] for inference in images of arbitrary sizes. For all inferences, the following set of sliced inference parameters was used:

- Overlap height and width = 0.2
- perform_standard_pred = False
- postprocess_type = “GREEDYNMM”
- postprocess_match_metric = “IOS”
- postprocess_match_threshold = 0.3

Maximal number of detections was explicitly increased to 100000 in sahi folder, models/ultralytics.py file.

To test how models perform on counting cells from the LIVECell dataset, we downloaded the dataset from https://sartoriusresearch.github.io/LIVECell/. We used pycocotools library to fetch the number of masks on each image. We also excluded images from LIVECell that have double sets of annotations. To test models, we combined train, validation and test images of each cell line into one set, then 10% of it was used to calibrate the model (calculate optimal confidence threshold), and 90% was used for the test. 410 pixels SAHI slice size was used. Mean Absolute Percentage Error (MAPE), among images of certain cell type and scatter plot of Predicted Counts vs. Ground Truth Counts were used as metrics of counting accuracy. MAPE was calculated as

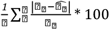

where ⍰ _⍰_ – ground truth number of cells on an image i taken from LIVECell annotations, 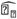 – number of cells predicted by a model for the same image. Scatterplots were generated using the seaborn 0.13.2 library.

To test how suitable models’ predictions are for single-cell tracking, we used Cell Tracking Challenge [16] datasets: “Glioblastoma-astrocytoma U373 cells on a polyacrylamide substrate” to test how models perform on phase contrast images of different cells, and “HeLa cells on a flat glass” to test how models perform not only on a different cell type, but on a different imaging modality. Datasets were downloaded from https://celltrackingchallenge.net/2d-datasets/. Using the Napari GUI and NuclePhaser prototype, we created two sets of tracking markers: unedited nuclei detection model’s predictions and manual annotations, which were used as ground truth markers. Then we created two sets of tracks from those markers with the btrack [17] plugin with the same parameters for each movie. We used two biology-motivated metrics from Ulman et al. [21]: Complete Tracks (CT), defined as “fraction of ground truth cell tracks that a given method is capable of reconstructing in their entirety, from the frame they appear in, to the frame they disappear from”, and Branching Correctness (BC), defined as number of divisions from ground truth also detected in the tested tracks; tracks derived from manual annotations were used as a ground truth. Cells that leave field of view during the movie were not used in the test. Tracks derived from predictions and ground truth tracks were compared manually.

