## Supplemental Table 1 for "NuclePhaser: a YOLO-based framework for cell nuclei detection and counting in phase contrast images of arbitrary size with support of fast calibration and testing on specific use cases"

Supplementary table 1. Results of validation on subsets of images of different cell types. Best corresponding metrics for each cell type are highlighted in bold.

| Model | Metric | CHO | HEK293 | iPSC | MSC |
|---|---|---|---|---|---|
| v5n | mAP 0.5 | 0.885 | 0.798 | 0.883 | 0.715 |
| | mAP 0.5-0.95 | 0.467 | 0.373 | 0.526 | 0.348 |
| | Precision at top F1 | 0.872 | 0.802 | 0.88 | 0.795 |
| | Recall at top F1 | 0.824 | 0.735 | 0.811 | 0.633 |
| | Top F1 | 0.847 | 0.767 | 0.844 | 0.705 |
| v5s | mAP 0.5 | 0.916 | 0.836 | 0.911 | 0.763 |
| | mAP 0.5-0.95 | 0.521 | 0.424 | 0.578 | 0.395 |
| | Precision at top F1 | 0.892 | 0.822 | 0.893 | 0.824 |
| | Recall at top F1 | 0.855 | 0.775 | 0.842 | 0.675 |
| | Top F1 | 0.873 | 0.798 | 0.867 | 0.742 |
| v5m | mAP 0.5 | 0.932 | 0.861 | 0.929 | 0.784 |
| | mAP 0.5-0.95 | 0.548 | 0.463 | 0.62 | 0.422 |
| | Precision at top F1 | 0.903 | 0.837 | 0.905 | 0.838 |
| | Recall at top F1 | 0.871 | 0.8 | 0.863 | 0.697 |
| | Top F1 | 0.887 | 0.818 | 0.884 | 0.761 |
| v5l | mAP 0.5 | **0.941** | 0.869 | **0.937** | **0.795** |
| | mAP 0.5-0.95 | 0.568 | 0.48 | 0.637 | 0.437 |
| | Precision at top F1 | **0.91** | **0.845** | **0.91** | **0.842** |
| | Recall at top F1 | **0.879** | **0.806** | **0.87** | **0.707** |
| | Top F1 | **0.894** | **0.825** | **0.890** | **0.769** |
| v5x | mAP 0.5 | 0.932 | 0.864 | 0.923 | 0.76 |
| | mAP 0.5-0.95 | 0.542 | 0.465 | 0.611 | 0.407 |
| | Precision at top F1 | 0.903 | 0.844 | 0.904 | 0.826 |
| | Recall at top F1 | 0.87 | 0.799 | 0.862 | 0.684 |
| | Top F1 | 0.886 | 0.821 | 0.883 | 0.748 |

| Model | Metric | CHO | HEK293 | iPSC | MSC |
|---|---|---|---|---|---|
| v11n | mAP 0.5 | 0.854 | 0.808 | 0.886 | 0.693 |
| | mAP 0.5-0.95 | 0.461 | 0.386 | 0.547 | 0.357 |
| | Precision at top F1 | 0.844 | 0.8 | 0.869 | 0.802 |
| | Recall at top F1 | 0.788 | 0.725 | 0.802 | 0.582 |
| | Top F1 | 0.815 | 0.761 | 0.834 | 0.675 |
| v11s | mAP 0.5 | 0.842 | 0.857 | 0.956 | 0.74 |
| | mAP 0.5-0.95 | 0.411 | 0.427 | 0.626 | 0.398 |
| | Precision at top F1 | 0.853 | 0.837 | 0.889 | 0.824 |
| | Recall at top F1 | 0.774 | 0.774 | 0.832 | 0.629 |
| | Top F1 | 0.812 | 0.804 | 0.860 | 0.713 |
| v11m | mAP 0.5 | 0.891 | 0.878 | 0.931 | 0.774 |
| | mAP 0.5-0.95 | 0.489 | 0.496 | 0.632 | 0.44 |
| | Precision at top F1 | 0.857 | 0.836 | 0.902 | 0.836 |
| | Recall at top F1 | 0.817 | 0.802 | 0.853 | 0.664 |
| | Top F1 | 0.837 | 0.819 | 0.877 | 0.740 |
| v11l | mAP 0.5 | 0.92 | **0.88** | 0.933 | 0.778 |
| | mAP 0.5-0.95 | 0.578 | 0.5 | 0.636 | 0.445 |
| | Precision at top F1 | 0.885 | 0.837 | 0.905 | 0.837 |
| | Recall at top F1 | 0.849 | 0.804 | 0.854 | 0.669 |
| | Top F1 | 0.867 | 0.820 | 0.879 | 0.744 |
| v11x | mAP 0.5 | 0.925 | **0.88** | 0.932 | 0.78 |
| | mAP 0.5-0.95 | **0.59** | **0.504** | **0.638** | **0.446** |
| | Precision at top F1 | 0.888 | 0.838 | 0.905 | 0.841 |
| | Recall at top F1 | 0.853 | 0.802 | 0.853 | 0.669 |
| | Top F1 | 0.870 | 0.820 | 0.878 | 0.745 |
