## Supplemental Table 2 for "NuclePhaser: a YOLO-based framework for cell nuclei detection and counting in phase contrast images of arbitrary size with support of fast calibration and testing on specific use cases"

Supplementary table 2. Results of count test of YOLO models on LIVECell dataset.

| Cells | Model | Optimal threshold | MAPE | | Cells | Model | Optimal threshold | MAPE |
|---|---|---|---|---|---|---|---|---|
| A172 | v5n | 0.21 | 11.0% | | MCF7 | v5n | 0.24 | 14.1% |
| | v5s | 0.22 | 10.7% | | | v5s | 0.29 | 13.0% |
| | v5m | 0.15 | 9.9% | | | v5m | 0.21 | 13.2% |
| | v5l | 0.27 | 10.2% | | | v5l | 0.4 | 13.5% |
| | v5x | 0.16 | 10.8% | | | v5x | 0.22 | 16.1% |
| | v11n | 0.1 | 10.6% | | | v11n | 0.08 | 18.6% |
| | v11s | 0.13 | 10.2% | | | v11s | 0.11 | 16.1% |
| | v11m | 0.09 | 12.7% | | | v11m | 0.11 | 19.0% |
| | v11l | 0.15 | 10.9% | | | v11l | 0.18 | 17.7% |
| | v11x | 0.2 | 10.2% | | | v11x | 0.22 | 13.4% |
| BT474 | v5n | 0.22 | 17.6% | | SHSY5Y | v5n | 0.17 | 15.2% |
| | v5s | 0.25 | 17.1% | | | v5s | 0.17 | 13.8% |
| | v5m | 0.07 | 27.6% | | | v5m | 0.13 | 17.6% |
| | v5l | 0.36 | 15.3% | | | v5l | 0.32 | 14.1% |
| | v5x | 0.09 | 14.9% | | | v5x | 0.08 | 21.2% |
| | v11n | 0.07 | 19.1% | | | v11n | 0.09 | 26.4% |
| | v11s | 0.06 | 19.1% | | | v11s | 0.08 | 26.4% |
| | v11m | 0.06 | 31.9% | | | v11m | 0.07 | 30.5% |
| | v11l | 0.05 | 25.2% | | | v11l | 0.1 | 18.1% |
| | v11x | 0.15 | 15.0% | | | v11x | 0.19 | 13.7% |
| BV2 | v5n | 0.29 | 14.2% | | SKBR3 | v5n | 0.24 | 3.2% |
| | v5s | 0.26 | 14.8% | | | v5s | 0.23 | 3.3% |
| | v5m | 0.25 | 23.2% | | | v5m | 0.21 | 4.1% |
| | v5l | 0.28 | 20.1% | | | v5l | 0.26 | 3.4% |
| | v5x | 0.19 | 19.4% | | | v5x | 0.13 | 3.7% |
| | v11n | 0.19 | 14.3% | | | v11n | 0.14 | 3.2% |
| | v11s | 0.24 | 15.0% | | | v11s | 0.15 | 3.1% |
| | v11m | 0.16 | 18.5% | | | v11m | 0.2 | 3.8% |

| | | | | | | | | |
|---|---|---|---|---|---|---|---|---|
| | v11l | 0.15 | 28.6% | | | v11l | 0.12 | 4.6% |
| | v11x | 0.14 | 17.7% | | | v11x | 0.17 | 3.5% |
| HUH7 | v5n | 0.3 | 17.6% | | SKOV3 | v5n | 0.15 | 6.4% |
| | v5s | 0.21 | 18.5% | | | v5s | 0.11 | 6.5% |
| | v5m | 0.09 | 18.5% | | | v5m | 0.09 | 6.8% |
| | v5l | 0.22 | 16.4% | | | v5l | 0.21 | 6.5% |
| | v5x | 0.17 | 14.9% | | | v5x | 0.07 | 6.0% |
| | v11n | 0.16 | 20.2% | | | v11n | 0.07 | 7.4% |
| | | | | | | v11s | 0.1 | 9.3% |
| | v11s | 0.13 | 18.9% | | | | | |
| | v11m | 0.07 | 32.4% | | | v11m | 0.08 | 9.2% |
| | v11l | 0.09 | 25.7% | | | v11l | 0.09 | 9.9% |
| | v11x | 0.19 | 18.7% | | | v11x | 0.11 | 7.8% |
